# Design and implementation of an asynchronous online course-based undergraduate research experience (CURE) in computational genomics

**DOI:** 10.1101/2023.11.29.569298

**Authors:** Seema Plaisier, Danielle O. Alarid, Sara E. Brownell, Kenneth Buetow, Katelyn M. Cooper, Melissa A. Wilson

## Abstract

As genomics and information technologies advance, there is a growing demand for research scientists trained in bioinformatics methods to determine gene expression underlying cell biology in health and disease. One approach to increase the number of scientists proficient in bioinformatics is to expand access through online degree programs and remotely-accessible learning materials. Fully-online learners represent a significant and growing community of historically underrepresented students who are frequently excluded from research opportunities that require in-person attendance during standard operational hours. To address this opportunity gap, we developed an asynchronous course-based undergraduate research experience (CURE) for computational genomics specifically for fully-online biology students. We generated custom learning materials and leveraged remotely-accessible resources on a high performance computing cluster to address a novel research question: the effect of changing quality trimming parameters for RNA sequencing reads on the discovery of sex-based differential gene expression in the human placenta. Here we present the process by which the instructional team devised and distributed analysis to address this question over a 7.5-week CURE and provided students with concurrent training in biology, statistics, computer programming, and professional development integral to the successful execution of the project and future publications. Scores from identical learning assessments administered before and after completion of the CURE showed significant learning gains across biology and coding course objectives. Open-response progress reports were submitted weekly and identified self-reported adaptive coping strategies for challenges encountered throughout the course. The instruction team monitored the progress reports to identify problems that could be resolved through collaboration with instructors and peers via messaging platforms and virtual meetings. Analytics from the course messaging platform demonstrated that high posting engagement was strongly correlated to high normalized learning gains, showing that students can effectively use asynchronous communication platforms to facilitate learning. The online genomics CURE resulted in unanticipated positive outcomes, including students voluntarily extending their participation beyond the course duration, presenting their findings at research symposiums, and applying to graduate school. These outcomes underscore the effectiveness of this genomics CURE for training and recruitment purposes and demonstrate that students can be successful in online STEM-based research experiences if given channels for communication, bespoke and accessible learning materials, and the support of experts in the field. Online CUREs can provide valuable research experience to harness the potential of online STEM students towards a more skilled, diverse, and inclusive workforce for the advancement of biomedical science.

## Introduction

Over the last twenty years, biomedical science has seen enormous growth in the amount of genomic data produced to investigate the molecular underpinnings of cell biology in health and disease. Resources requiring-omics (e.g., transcriptomics, genomics, proteomics) level analyses are often hosted on high-performance biocomputing clusters or cloud computing environments by which users may log in from any location as long as they have a computer and sufficient internet access. As high-throughput molecular assays and technology for data processing and machine learning advance, there is an increasing need for cross-disciplinary computational analysts trained to understand biology, genetics, statistics, pharmacology, and mathematical modeling. Students who develop skills in computation and programming are more prepared to answer biological research questions and benefit from the high demand of these skills in the STEM industry (Wilson Sayres et al. 2018; Bennett 2020; Gao and Guo 2023). Thus, building a computational genomics workforce is a necessary step to maximize the knowledge gained from the outpouring of molecular profiling data and computational genomics research is uniquely suited for asynchronous online learning.

Large scale survey data has shown that undergraduate research experiences increase interest in STEM careers, gains in research skills, and the likelihood that students will pursue graduate degrees (Russell, Hancock, and McCullough 2007; Lopatto 2004, 2007; Paalman 2002). While these historically have taken the form of apprenticeships where students work in a faculty member’s lab, there are too few research opportunities for all undergraduate biology students to be able to participate (Bangera and Brownell 2014). An alternative approach is to broaden access to research by developing course-based undergraduate research experiences (CUREs). CUREs are formal courses in which students use well-established scientific practices to participate in novel research projects with undetermined outcomes of interest to the broader scientific community (Auchincloss et al. 2014). CUREs can be effectively developed using a “backwards design” in which a research question and appropriate experimental plan are determined and then learning objectives of the course are derived to fit that research plan (Cooper, Soneral, and Brownell 2017). CURE topics are often driven by questions that arise in the instructing faculty’s area of interest, thereby providing students with mentors and skills as well as providing mentors with a pipeline to train and recruit students that can advance research programs (Shortlidge, Bangera, and Brownell 2015). CURE research projects are aimed at publication, which allows both mentors and students to contribute to the field while allowing the students to gain key research skills and build a science identity (Rubenstein et al. 2023).

Students who participate in CUREs can be familiarized with the cultural norms of scientific research, learn to cope with scientific challenges, and be better prepared for future research projects and graduate school (Bangera and Brownell 2014; Bennett 2020; Gin et al. 2018). Fully-online students have shown the same amount of interest in conducting research as their on-campus peers, but have reported a significantly lower awareness of accessible opportunities (Faulconer et al. 2020; Cooper, Gin, and Brownell 2019). This may be attributed to the fact that most research experiences, including CUREs, published up to the time of this study were formatted for in-person or hybrid experiences rather than for asynchronous, online students. However, there were some reported exceptions that transitioned to online and remote learning in response to the COVID-19 pandemic (DeHaven et al. 2022). Increasing and embedding online research into course schedules can expand access to these learning benefits and continuously improve research programs to adapt to the growing population of online learners.

In addition to limited online CURE offerings, asynchronous students are often confined by their inability to attend programs in person due to geographic location, work schedules, and life obligations (among other responsibilities), yet have demonstrated the same ability to make learning gains in online biology courses (Paul and Jefferson 2019). Additionally, online undergraduates are more likely to be from underrepresented demographics including first generation college students, women, low-income households, and non-traditional adult learners (Bangera and Brownell 2014; Mead et al. 2020). As such, restricting CURE offerings to on-campus facilities only adds another barrier for online students to overcome. Online research experiences have the potential to increase the amount of historically-underrepresented student participation in STEM, increase their application to (and likely enrollment into) graduate programs, and open additional career opportunities, thereby making STEM education more equitable and the future workforce more diverse (Cooper, Gin, and Brownell 2019; Merrell et al. 2022; Faulconer et al. 2020).

Bioinformatics represents a unique opportunity for the development of fully online CUREs. The research can be conducted with a computer and internet access, making it highly accessible. Second, student anxieties around genomics and bioinformatics can be lessened through the collaborative CURE community, which is typically a larger community than most undergraduates doing research have. Engaging with bioinformatics in a CURE setting can give students more detailed instruction on how to address common sources of anxiety, which for bioinformatics research can include a lack of pertinent background knowledge, computer programming anxiety and inexperience, and issues related to accessibility and inclusion in the virtual classroom (Gin et al. 2018; Forrester et al. 2022). Worry or nervousness about conducting research itself can be stressful for students, particularly when the students are from underrepresented or underserved communities (Cooper, Eddy, and Brownell 2023; Pennino et al. 2022). While these factors have been linked to high attrition rates in online STEM courses, the literature shows that retention can increase with student-specific interventions (Wladis, Hachey, and Conway 2014). In this course, the instruction team leveraged the asynchronous capabilities of institutionally-approved messaging platforms (Slack) and encouraged open-dialog in recorded, virtual lab meetings and weekly progress reports where students self-identified the strategies acquired to cope with the challenges of online research. Generally, coping strategies can be adaptive (active) such as seeking support, planning, or taking specific actions to solve problems, or maladaptive (evasive) such as distraction or denial (Skinner et al. 2003); encouraging students to identify and engage in effective coping strategies can help them develop proactive habits that motivate them to commit to STEM courses, degrees, and employment opportunities (Major, Holland, and Oborn 2012). Since this research project was novel and bespoke code was run with course datasets for the first time, the instruction team closely monitored student submissions and Slack posts, and were able to intervene with individuals who were transparent with the challenges they experienced and the coping strategies they employed. Because students were being guided through the research process in a supportive, collaborative, and iterative manner, maladaptive coping strategies were addressed individually and adaptive coping strategies were shared with the class then noted for course improvement in future iterations.

In this manuscript, we describe how we developed and implemented an online CURE in computational genomics to study a specific part of data preprocessing for sex-based differential gene expression analysis. We discuss how the required analysis was distributed among students in an asynchronous format over 7.5 week-long modules and how research-specific learning materials were designed to promote student success. We demonstrate that our implementation of this CURE led to improvements in total assessment scores as well as specifically within the biology/statistics, coding, and professional development sections. We describe the coping strategies students used to address challenges encountered while conducting bioinformatics research and how support was given in an asynchronous learning environment. As guidance on running an asynchronous CURE is challenging to find, this discussion on how a computational genomics CURE was designed for online students is intended to serve as a template for others seeking to expand inclusive and accessible research opportunities at their educational institutions.

## Course Design and Implementation

### Research project

The research project at the center of this course investigated the effect of data processing methods on identification of sex differences in gene expression in human placenta. In a previously published study (Olney et al. 2022), the instructors of this CURE identified genes that were differentially expressed between male and female placentas from full-term pregnancies using a well-established analysis workflow: trimming of low quality RNA sequencing reads and the limma-voom pipeline for differentially expression of genes between two groups (males and females; https://github.com/SexChrLab/Placenta_Sex_Diff). The parameters chosen for trimming adapter sequences and low-quality RNA sequencing reads were determined using experiences of the lead investigators of the study, but literature review showed a variety of thresholds for what was considered to be low quality for reads. This led to the research question of how robust the published sex differential gene expression profile was to changes in trimming parameters. Literature on the effect of changing thresholds for trimming on differential gene expression is both limited and contradictory (Williams et al. 2016; Liao and Shi 2020; Macmanes 2014; Del Fabbro et al. 2013), so the instructors devised a course that would allow students to test a range of values for the ‘trimq’ and ‘minlen’ trimming parameters implemented in bbduk, the software used in the original manuscript, to test whether and how the sex differential expression profile was affected in the placenta tissues. All of the required background knowledge for this project and project aims were presented to students in the first learning module of the course.

Sample data for this course was previously published RNA sequencing data from placenta samples taken at full-term deliveries (Olney et al. 2022). Only the first of two batches of samples collected was used to avoid the need to correct for batch effect, giving 10 placentas from female-carrying pregnancies and 12 placentas from male-carrying pregnancies. Each placenta was sampled twice and each sample was processed for bulk RNA sequencing; gene expression for samples from the same placenta was summed.

### Course Timeline

The asynchronous genomics CURE was offered for 3 credits and ran for a 7.5-week session (Fall 2022 B session) after a required prerequisite 7.5 week course in the first half of the semester (BIO 439: Computing for Research; Fall 2022 A session). Each module was one week long; students completed the reading and assignments while instructors maintained an active presence on Slack to offer support and collected weekly progress reports (**Figure 1**). In the prerequisite course, which assumed no prior coding experience, students were introduced to command line programming, bash scripting, navigating a high performance computing cluster (hpc), and ran through basic genomics analysis (fastq file quality control, alignment, and variant calling). The CURE was organized into seven modules, each focusing on three areas: Biology/Statistics, Coding, and Professional Development. Time estimates were provided separately for each section of each learning module to help students plan their work time.

**Figure 1:**
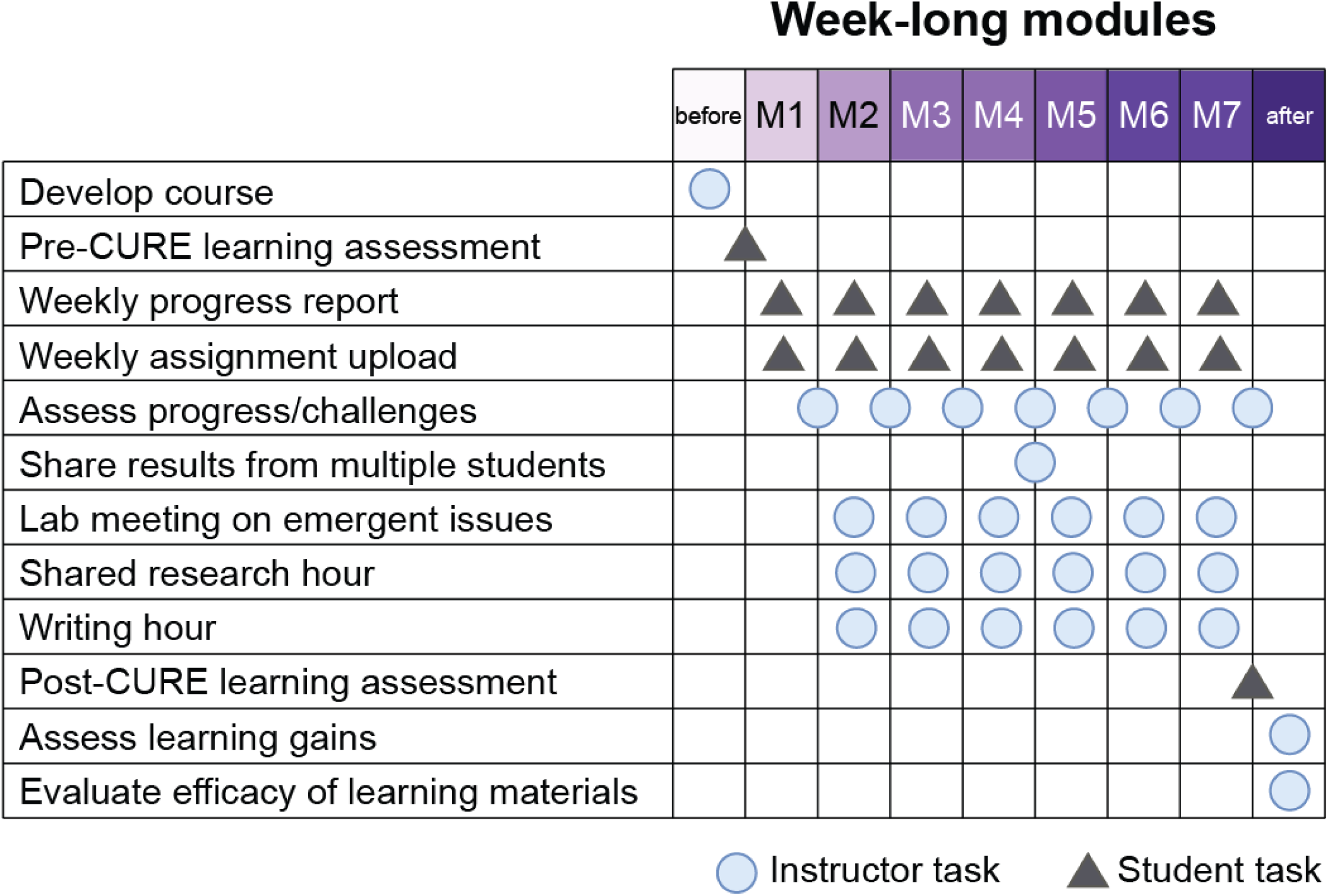
Instructor and student contributions over 7-week CURE. Instructor contributions are labeled as circles, student contributions in triangles. Students began by filling out a pre-assessment to show their baseline level of knowledge on the research topic. Each week students completed research goals and turned them in as assignments along with a progress report communicating their achievements and challenges. Instructors used both of these to select topics for weekly lab meetings using teleconference software and recorded for students that could not attend. Instructors hosted shared research (office) hours throughout the week. Instructors monitored progress on research goals and distributed data generated by individual students at the midpoint of the CURE. Following the completion of the course, students completed a post-assessment survey and instructors analyzed which areas students were able to increase their knowledge and potential misconceptions that can be addressed after the course and/or in future CUREs.

### Intended audience and prerequisite knowledge

The CURE was open to students who completed the prerequisite course (BIO 439: Computing for Research) and were enrolled through Arizona State University Online. This is a fully online course with no in-person components. The prerequisite course introduced the students to using genomics software tools in a Linux/Unix shell environment to learn, apply, and write code to process high-throughput sequencing data. Initial enrollment for the CURE was capped at 20 students, of which 15 total students completed the course and 13 completed both the pre– and post-assessments and informed consent to have their learning assessments analyzed.

### Distribution of analysis among students

Once the research project and course schedule had been defined, a plan to distribute the work among the enrolled students was devised to fit in the 7 week-long modules allotted for the course (**Figure 2**). We designed the analysis steps such that all students were working on the same analysis at the beginning of the course, so that they could ask each other for help and the instructors were clear on what results the students would see. For our research project, we performed data processing before the course started so that they could spend the first two modules learning the required conceptual background material to understand the research project and learn basic programming techniques needed to run differential expression analysis and data visualization. In Module 3, all students were assigned to run the differential gene expression pipeline on the raw data with no quality trimming applied. In doing this, they were able to learn how to use the code, modify it where necessary, and ask questions from their instructors and peers if they ran into any problems. For Module 4, we assigned data trimmed with specific trimming parameters to pairs of students to analyze using the same code they worked with all together with the untrimmed data. We paired the students so that they would have a way to double check that someone else got the same results they did and could have someone to work with if they desired or could just as easily work independently. The decision about whether to work in a group or individually was up to the student, and intended as an inclusive teaching strategy given time zone differences and the difficulty of doing group work asynchronously. In this way, we distributed the analysis equally among the students and simultaneously built in redundancy to make the research goals more robust to students having problems or dropping out of the course.

**Figure 2:**
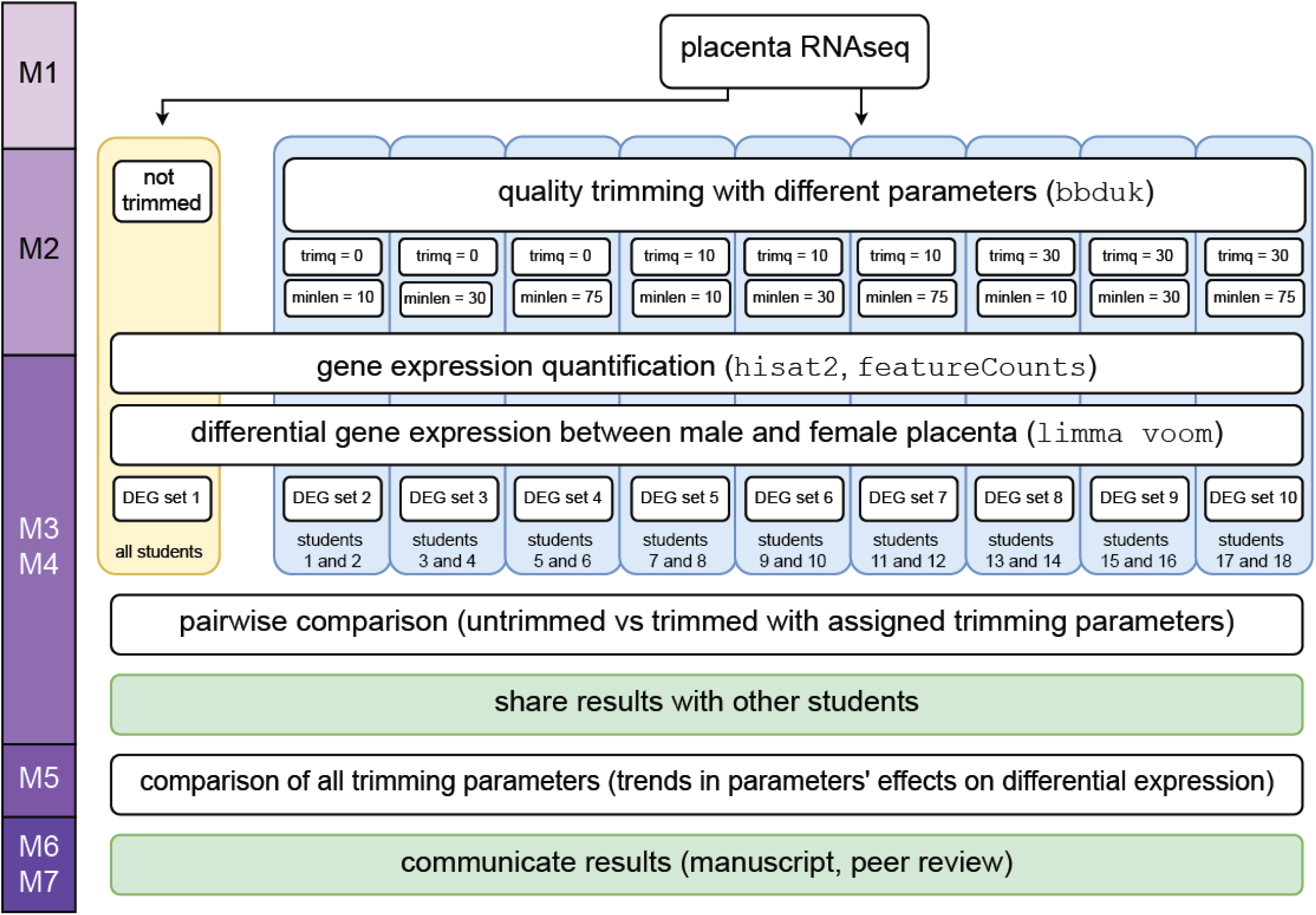
Distribution of analysis among students. To address the research question of how quality trimming of RNA sequencing data affects identification of genes differentially expressed by sex in the human placenta, data sets were created using 9 pairs of values for two parameters of trimming software (“trimq”: minimum trimming quality score, “minlen”: minimum length after trimming) to compare to the reads with no trimming applied. Each data set was preprocessed for gene expression quantification and divided among the students to identify sex differentially expressed genes. Results from students were shared so students can analyze trends across the full range of trimming parameters. Each student described the results of the study in a manuscript which was peer reviewed by other students.

Once the students had performed differential expression analysis, we had them each compare their results to those attained with the untrimmed data. At the end of Module 4, we asked that they turn in their code, differential gene expression profile, and results of the two-way comparison as a graded assignment. The instructors then checked for errors and put all the results in a central online location so that the entire class could access it. Then for Module 5, the students were able to use all the results generated by the entire class to compare results across the full trimming parameter range to determine trends in how trimming parameters affected differential gene expression. With access to the fully analyzed data set, each student was asked to present results in the format of a scientific manuscript and peer review manuscripts written by classmates.

### Student instructions

To set the expectations for the students, the lead instructor began the course by recording an introductory video where she stated that the CURE would guide students through working on a real research project and that the instructors did not know what the outcome would be in advance. She indicated that the students are being graded on their problem solving and their own dedication to advancing the research goals, not the specific outcome of the analysis. She indicated that it was acceptable for the students to feel challenged for any reason (inexperience with research, programming, newness to the field, etc.), but that the students should reach out to the instructors and other students in the class and take advantage of their resources to address those challenges. Students were encouraged to ask questions on the direct messaging platform Slack so that all students in the class could view the responses and learn together. Students were told that the research project would be continued after the CURE and that there would be opportunities for authorship in future symposium presentations and publications.

Students were shown a variety of resources that they could use during and after the CURE. Each student was given an account to the university’s high-performance biocomputing cluster free-of-charge; access could be granted after the CURE if the student continued with the project with the lead instructor as a mentor. They were pointed to various resources for learning how to use the cluster as well as ways to ask the research computing department for help. The CURE was maintained using Canvas, the same education management system that was used throughout the institution for accessing grades, syllabi, etc., so that students would feel comfortable with submitting assignments and receiving feedback during assignment grading.

In addition to the research project being conducted, the students were told that their participation in the CURE helps to shape future CUREs. By filling out the pre– and post-assessments and the study consent forms, the students were part of an educational study where the instructors could assess what teaching methods gave rise to the most student learning and which concepts were most likely to harbor misconceptions. This explanation was intended to increase the students’ investment into their participation in a CURE and increase their communication and engagement.

### Faculty instructions

Expectations of the lead faculty instructor were to work with the other course instructors on developing a research project of interest to the field. All course instructors were expected to be knowledgeable about the research project and guide the students through interpreting and contextualizing their findings, but not expected to know the answer to every question any student asked. The nature of a CURE being based on a novel research question meant that there would be questions that no one knew the answer to. Instructors were expected to help troubleshoot code students developed for the research project and use their expertise to show students how to read related publications or interrogate the data further. Instructors were expected to grade assignments and progress reports with predetermined rubrics as a way of monitoring progress on the research project.

## Course Materials and Methods

### Learning objectives

Once the overall experimental design plan was determined, custom learning objectives were written to outline assessable skills students would gain during the completion of the research project (**Table 1**). Learning objectives and corresponding learning materials were written in a modular fashion such that blocks could be reused for future CUREs based on what is needed for the specific research question being investigated by the instructors and students. Certain objectives were specific to this research project regarding the effect of trimming on sex differential expression, but others were about general topics which could be reused for any other CURE research projects that involve that topic. Instructors basing projects on their own laboratory’s research focus will likely find they can build a set of topics that will be in common for many new projects that are devised, thus reducing the amount of time needed to develop learning resources for future CUREs and formally creating training materials for personnel entering their research group. Furthermore, professional development blocks are general skills that are important across all research projects.

**Table 1.** Genomics CURE learning objectives by module and submodule. Learning objectives designed around the specific aims of the CURE were divided into 7 week-long modules (rows) and further subdivided into three submodules addressing the Biology/Statistics, Coding, and Professional Development skills gained as the students conducted research (columns). Objectives colored in orange are specific to each genomics CURE project, so learning materials for these sorts of objectives will have to be newly generated for each future genomics CURE. Objectives colored in purple form the foundation of any differential gene expression experiment, so learning materials for these objectives can be reused for any future genomics CURE that involves differential gene expression. Objectives colored in green are essential skills for any bioinformatics researcher, so learning materials for these objectives can be reused for any future bioinformatics CURE project.

| Learning Objective | Biology | Coding | Prof. Dev. |
| --- | --- | --- | --- |
| 1. Understand the research project | Understand research aims and experimental design<br>Describe function of placenta and sex chromosomes<br>Understand next-generation sequencing methods | Navigate files/directories in Unix shell<br>Connect to the RStudio server | Use Google to search for useful R solutions<br>Practice reading a scientific article |
| 2. Quantify and normalize gene expression | Understand how a differential expression experiment is designed and how results are analyzed<br>Understand trimming software | Run differential expression pipeline in R | Search for R packages<br>Read a relevant scientific article |
| 3. Identify differentially expressed genes (DEG) | Describe how linear modeling can be used to identify DEG<br>Describe how to visualize samples based on gene expression<br>Normalize RNA seq data<br>Describe effects of trimming | Visualize sample groupings with MDS plot<br>Visualize differential gene expression with box plots, violin plots, jitter plots, or a combination | Solve coding errors |
| 4. Compare two DEG lists | Overlap between two conditions | Visualize overlap between two groups with Venn diagrams<br>Visualize correlation with a scatter plot | Report details of analysis methods to ensure reproducibility |
| 5. Compare full range (>2) of DEG lists | Overlap analysis for higher number of comparisons | Visualize overlap between greater than 2 groups with Upset plots | Write a scientific paper |
| 6. Write a research paper | Understand how to cite publications<br>Write figure legends | Write the methods section of paper | Understand how authorship is |
|  | Write sections of paper that describe the study and the results |  | determined for publications<br>Understand ethics of data sharing |
| 7. Conduct peer review | Constructively review 3 papers | Submit all code, output files, and reports with descriptions to be stored/ archived | Propose relevant follow-up questions for future research |

### Module Learning Pages

For the text of each lesson, in lieu of expensive textbooks, the instruction team wrote collaboratively to produce bespoke text and reference materials to guide the students as they performed the analysis for the research project. Each module had web pages posted in the Canvas learning management system in advance and was available for the students to reference throughout the course as needed. Each learning page had three sections based on the learning objectives: Biology/Statistics, Coding, and Professional Development. The instruction team surveyed publicly available videos, primary literature, and publicly available descriptions and created custom text, figures, and videos to create background materials specific to the chosen research project. To minimize cognitive load and maintain subject relevancy, videos were generally kept to under five minutes long. Shorter videos focused on specific topics make it possible to arrange or update specific topics to customize learning materials for future CUREs. Additional learning resources for novices and enrichment for more advanced students were included as “Additional Resources”. Published bioinformatics workflows, vignettes, and tutorials were incorporated and/or modified as needed to demonstrate analysis relevant to answering the research question.

The instructional team aimed to increase accessibility and inclusion when designing learning materials for each module. Communication considerations were at the forefront of the course design and asynchronously leveraged direct messaging platforms (Slack), coordinated video conferences with transcriptions and recordings available, and Canvas-integrated tools such as announcements and email. All videos were downloaded without advertisements and fully transcribed by the instruction team to provide students with a text alternative. Learning pages and software were at no added cost to students as project-specific materials were authored by the instruction team and from publicly-available resources. To reduce the need for expensive personal computers, students were strongly encouraged to use the RStudio Server offered by the institution’s high performance computing cluster which reduces the need to store large data files and bulky software packages on local drives. Programming was done with RStudio, a freely available, cross-platform coding environment which the students can continue using through the institution’s high-performance biocomputing cluster or download locally for use after the course was complete.

## Student assessments

### Pre– and post– assessments

Students took an identical learning assessment before and after completing the genomics CURE. Questions were designed by the course instructors and vetted by outside genomics experts to match the learning objectives of the course. Scores and answers were not revealed to the students to remove any advantages for the post-assessment. The intent behind the learning pre-assessment was to provide a baseline of student subject knowledge from which gain was measured after course completion. The learning pre– and post-assessments had a total of 25 questions with seven addressing biology/statistics objectives, seven addressing coding, four addressing professional development, and seven Likert surveys addressing student comfort levels in computational research. Students received one attempt to complete the pre-assessment and would receive full credit for the assignment regardless of score. Questions written in check-all-that-apply, matching, and multiple-choice for Likert survey formats. To ensure students were getting the correct answers for understanding specific concepts, the check-all-that-apply and matching questions had varying amounts of distractors and answers to be selected, ranging in five to six total options for most questions. To reduce anxiety levels, there was no time limit on test completion and no video proctoring (Gin et al. 2022), however, a submission was required to move on to subsequent modules.

### Weekly progress reports

To maintain an authentic research experience, the instruction team opted to develop an open-response summative assessment for students to submit weekly, designed in the fashion of progress reports submitted in the lead faculty instructor’s laboratory. To reduce student grading anxiety and to treat the assignment as an authentic update, report grading was flexible and geared towards completion over correctness. The report prompted students to list their successes in learned and applied content, perceived and technical challenges, how they addressed those challenges, and their plans for progressing through subsequent modules (**Table 2**).

**Table 2.** Genomics CURE Weekly Progress Report prompts. The categories of the open-response Weekly Progress Report include successes, challenges, how those challenges were addressed (coping strategies), and upcoming goals. Exploring student challenges and coping strategies in an open response format was essential for instructors to modify weekly meeting topics as needed. As mentioned, to maintain the integrity of the novel research, the instruction team did not run the full course pipeline, and the weekly progress reports became fundamental for keeping the pulse of class progress and to identify any major troubleshooting needed.

| Category | Prompt |
| --- | --- |
| Accomplishments | • List novel findings<br>• Concepts learned<br>• Coding solutions<br>• List successful communication with instructors and classmates |
| Challenges and how you addressed them | • List specific challenges for the week<br>◦ If you did not have challenges, describe your strategies/background used to make this a challenge-free week<br>• List your approaches for addressing this challenge (and if it is still outstanding)<br>◦ If you did not have challenges, describe how you helped others address a challenge (via Slack, meetings, group discussion) |
| Goals for the next week | • List how current module's objectives relate to next module<br>• List goals of the next module<br>• List content you want to revisit, or you think would be valuable to learn more about |

Exploring student challenges and coping strategies in an open response format was essential for instructors to modify weekly meeting topics as needed. As mentioned, to maintain the integrity of the novel research, the instruction team did not complete the analysis the students were meant to complete in advance, and the weekly progress reports became fundamental for keeping a pulse on the progress of the class and to identify any major troubleshooting needs.

### Coding assignments

Research milestones were implemented using weekly coding assignments, typically requiring upload of code reports and data output files. In order to help bridge the gap in coding experience between students in the class, the instruction team developed bespoke code which the students could modify and build on to complete the analysis required for the research project. Template code was provided in RMarkdown format so that code, products of code, and description of what the code was doing could be easily created for submission, communication with others, and documentation of methods at publication stage. Template code was highly commented to explain each computational step and encourage best practices for coding techniques.

Coding assignments were used to monitor the students progress through the parts of the research project and give feedback to students if they had problems. Each assignment had a custom rubric designed to make sure the students were understanding the key concepts being detailed in each module before moving on to the next step in the research plan. Because more value was placed on effort and progress than on specific outcomes, coding assignments were only worth 25 points while progress reports were worth 100 points in the total points for the course final grade.

### Manuscript and peer review

As a final project in the CURE, students were asked to write a report in the format of a manuscript for publication and then asked to conduct peer reviews on 3 manuscripts written by their classmates. A manuscript was chosen as the final assignment in order to formally teach students how to communicate scientific results with the level of accuracy and detail expected at a professional level. The professional development section of several modules was used to walk students through how to put together the various parts of their manuscript. Early modules showed students the sections of a scientific manuscript and demonstrated the type of information that goes into each section using published papers that served as background to the research project as examples. Middle modules presented the students with information and examples of how to write detailed legends for figures produced as the research project was conducted. Examples of how to keep track of and document software packages used in code in the methods section of the manuscript were featured in all coding templates. In this way, the students were able to collect relevant information from their assignments to begin writing their manuscripts.

The professional development section of Module 5 walks the student through how to storyboard the results attained throughout the CURE. A storyboard includes an outline of the aims of the research project, which topics need to be included in the introduction and methods, and an ordered list of figures with what results they felt they needed to include to state the overall results of the project. Storyboarding helped the students to understand how pieces of the analysis work together to produce an overall conclusion. It also helped the students to separate what was included in the course as background learning from what was done as part of the experimental design and research plan.

In Module 7, students were able to assess other students’ writing when asked to peer review 3 other students’ manuscripts. As all students did similar analysis even though the data they started with was different, students were able to appreciate the nuanced differences in how results were interpreted and explained. The rubric used to grade peer reviews of manuscript is provided (**Table 3**).

**Table 3.** Rubric for conducting peer reviews on student manuscripts.

| Direction | Prompt |
| --- | --- |
| For each review you are assigned, please comment on the following: | • Are there any points or sentences that are confusing to you?<br>• Do you feel there are places where the author needs more detail in order to make their point?<br>• Are the figures clearly labeled in the figure legend and clearly explained in the results?<br>• Can you follow the logic the author used to address their topic? Can you suggest an order for figures/results that you think would be more logical or give a better flow?<br>• Are there any statements that you feel need references?<br>• Are there any typos, spelling and grammar mistakes, or phrasing issues that you can point out to the author?<br>• Are there any parts of the paper that you feel are particularly awesome? Go ahead and bring attention to that with a reason you think it's awesome. |
| Please make sure to keep the following in mind for language: | • Keep your comments positive and helpful; remember that we are all in this together<br>• Do not suggest things that are out of scope or that you know would take too much time to do<br>• Read the entire report once all the way through before making comments so you know whether something you think is missing is just in a different order<br>• Try to be as specific as possible |

### Class Communication

Students were given several opportunities each week to meet with the instruction team in real time: lab meeting, writing hours, and shared research hours. The class was initially polled for availability to help choose times where the most students could attend real time meetings. All meetings were conducted over Zoom, allowing participants from any location around the world and live transcription to increase accessibility. Optional weekly lab meetings covered student-reported challenge topics and were recorded, transcribed, and posted with an announcement each week. Optional weekly writing hours gave students the opportunity to discuss weekly assignments, interpretation of findings, edit descriptions of the findings for assignments and their manuscript, and generally to refocus if they had challenges going into the next module. Optional shared research hours, similar to office hours, were offered on a biweekly basis, giving students the opportunity to troubleshoot code with the instruction team live with options to share their screen. Shared research hours were offered in the evenings and weekends to accommodate students that could not attend at traditional work hours.

Instructors frequently posted through the direct messaging platform Slack to ask about challenges, point out common errors, and identify opportunities for support. When the students were enrolled into the CURE, they were invited to a general Slack channel including all of the students enrolled and all of the instructors. Students were allowed to create direct message groups as needed, such as with the student that was assigned the same trimming parameter data set. Announcements made on Slack were also sent by email, but Slack allowed students to get fast responses from other students and instructors. Some troubleshooting issues were more appropriate for the institution’s research computing staff; in those cases, students were directed to send their Slack message to the research computing help desk Slack channel. Tips for posting code with proper formatting and sufficient background information were included in the professional development section of the learning pages, along with tips on how to effectively search for coding solutions with internet search engines.

For troubleshooting code, the interactive communication structure allowed the instruction team to discover areas of the code that were incorrect or needed debugging and to support students as they tried different coding solutions and battled the frustration of being novice programmers. If there were major troubleshooting issues discussed in Shared Research Hours or in Slack, the instruction team documented the steps taken to address problems including how the problems were identified and different solutions that were attempted before discovering the true solution. These documents were then distributed to students with announcements made in communication platforms and by email, in addition to presenting the information in the recorded and transcribed lab meetings.

## Results and Discussion

### Student learning gains

To determine if and what the students learned from the CURE, we used an identical learning assessment taken before and after the course. The results from the pre– and post-assessment included only first attempts at the examination and students who completed both assessments were evaluated for learning gains. Student pre-assessment and post-assessment results were exported from the Canvas learning management system (LMS) where they were administered and graded. The LMS only issued full credit for check-all-that-apply questions that had every selection correct, and deducted points for any distractor selection. For the purpose of this analysis, each question was worth 1.00-point total, and scores ranged depending on the number of options, correct answers, and distractors available per question. The eighteen total questions on biology, coding, and professional development learning outcomes were evaluated independently from the seven Likert survey questions.

To investigate the change in student performance between the pre– and post-assessments, we first compared the distribution of overall scores and per section scores before and after the CURE. A significant improvement in overall scores was observed with the mean score increasing by 11.64%, from 65.96% (SD = ±12.92) to 77.61% (SD = ±16.77, p = 0.003) (**Figure 3A**). This is considered to be a medium to large effect size as evaluated by Cohen’s d-statistic (d = 0.79, medium effect size range = 0.5-0.8, large effect size range > 0.8). There were significant increases in average Biology/Statistics scores (p = 0.0005) and Coding scores (p = 0.03). Students generally had high scores on Professional Development questions in the pre-assessment indicating that they came in with some familiarity with research papers and problem solving while doing research. Only a slight increase was observed in Professional Development questions in the post-assessment scores (p = 0.10) (**Figure 3B**). This demonstrates that it is indeed possible to achieve student learning gains in an asynchronous online undergraduate course.

**Figure 3.**
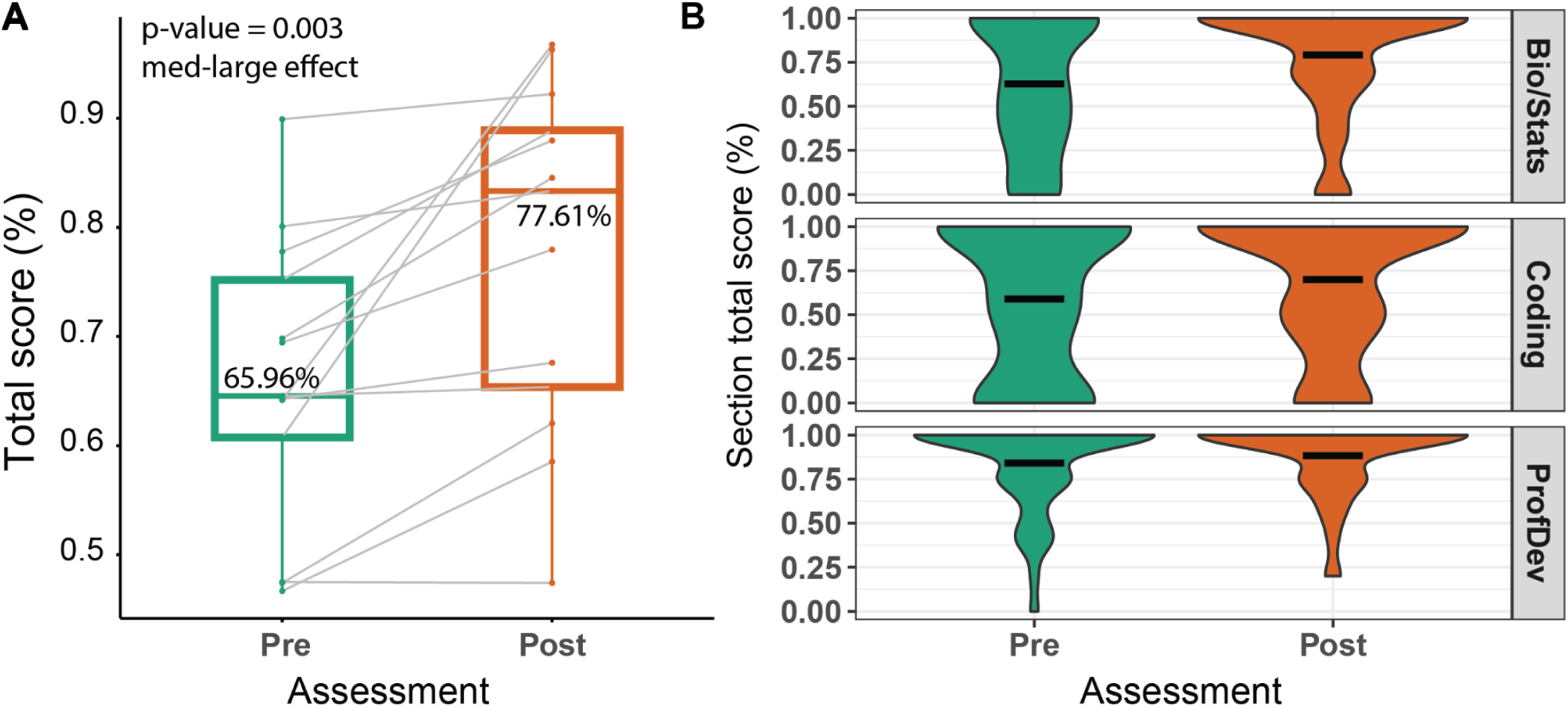
Student knowledge assessment before and after the course. (A) Boxplots depicting mean student assessment scores before (green) and after (orange) completing the Genomics CURE. Each point represents one of the thirteen students who completed both the pre– and post-assessments and the lines connect to the same student’s pre-assessment to post-assessment scores. The mean class score significantly increased by 11.64%, from 65.96% (SD = ±12.92) to 77.61% (SD = ±16.77, p = 0.003), an effect size considered to be a medium-large by Cohen’s d-statistc. (B) Violin plots depicting each pre-assessment (green) and post-assessment (orange) scores for all questions divided by topic: Biology/Statistics (7 questions), Coding (7 questions), and Professional Development (4 questions). Varying thicknesses of each plot represent the distribution of scores and widths between each peak represent score densities, or clusters of score occurrences.

Given that the students had a range of experience with coding, biology/statistics, and research, we assessed performance in the pre– and post-assessment as a proportion of the maximum possible increase in scores. For each question on the assessments, normalized learning gains (NLGs) were calculated as the difference between the post-assessment score and the pre-assessment score divided by the maximum potential gain from the pre-assessment score (DeHaven et al. 2022; Paustian et al. 2017; Colt et al. 2011) (**Figure 4**). Mean scores of each question were generally higher in the post-assessment, with positive normalized learning gains (NLGs) in 67% of questions (Q1, Q2, Q3, Q5, Q6, Q7, Q10, Q11, Q12, Q13, Q15, Q17), negative NLGs in 11% (Q4, Q8), and no changes in 22% (Q9, Q14, Q16, Q18) (**Figure 4**). For the two questions that had a negative learning gain, further analysis of Q4 demonstrated that answers chosen incorrectly in the post-assessment were likely the result of a misconception some of the students developed and answers chosen incorrectly for Q8 were due to confusion about the wording of the question. Mostly positive normalized learning gains demonstrated that the CURE allowed students to add to the knowledge they had before the CURE. Administration of a pre– and post-assessment provided insight into which aspects of the CURE were effective in providing learning gains and which led to misconceptions or otherwise needed improvement.

**Figure 4.**
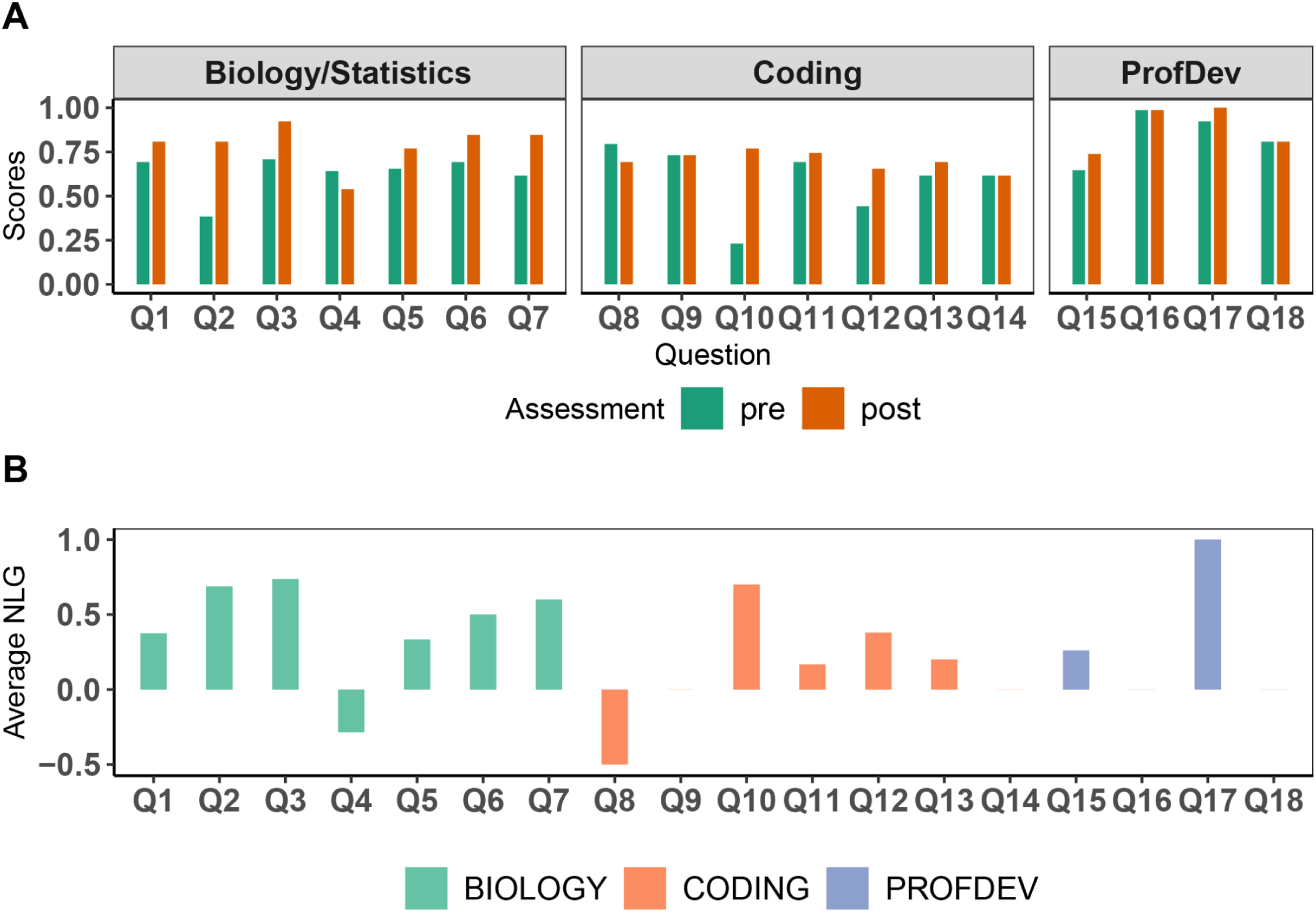
(A) Average question pre– and post-assessment scores divided by topic and (B) difference in normalized learning gain. A. Barplot depicting mean pre-assessment (green) and post-assessment (orange) scores divided by submodule learning topics. Most questions exhibited gains except for Q4 and Q8 which had decreases in mean scores.Q14, Q16, and Q18 had no mean score changes between assessments. B. Barplot depicting the normalized learning gain (NLG) per question, categorized by topic: biology (green), coding (orange), and professional development (purple). NLGs represent the magnitude of average learning gain per question and are determined by dividing the difference between post-assessment and pre-assessment scores by pre-assessment scores.

### Student coping strategies

The asynchronous CURE not only allows students from historically underrepresented backgrounds obtain research experiences, but also provides a chance for students to experience challenges endured in authentic research and allows them to identify coping strategies for these challenges in an environment where mistakes and failures come without severe penalty (Gin et al. 2018). Transitioning into computational research can be especially challenging for students, perhaps even more so in an asynchronous setting (Forrester et al. 2022; Joanne M McInnerney and Tim S Roberts 2004). Open-response weekly progress reports allowed the study of coping strategies employed by students throughout the CURE. Coping refers to the way students respond to extrinsic stressors that could increase anxiety and affect mental health. Coping strategies can be adaptive or maladaptive depending on whether they resolve stressors or prevent resolution respectively and are composed of different themes (Skinner et al. 2003; Musgrove et al. 2021; Henry et al. 2019).

Each week the instruction team would review challenges reported and how the student reacted to that challenge in the organic responses in the weekly progress reports. Students were prompted to list the coping strategies they used to address challenges, or, in cases of no reported challenges, the background knowledge or previous experience they leveraged to be successful. The instruction team synthesized and quantified the individual responses into a semantics bank to identify the top reported challenges and successes (**Figure 5**). Challenges were clustered into categories: biology and statistical analysis, coding, professional development, cognitive load, personal, and other. Five strategies were adaptive, including problem solving, support seeking, information seeking, self-reliance, and cognitive restructuring. Accommodation, negotiation, and distraction can be adaptive or maladaptive depending on how the student responded to the stressor, and strategies that are generally maladaptive include escape, rumination, helplessness, delegation, and opposition (Skinner et al. 2003). One instructor did the assignment of statements from the progress reports to coping strategy categories so that the assignments remained consistent throughout the course. Results indicated that adaptive coping strategies were employed throughout the CURE by the students (with problem solving reported at an especially high level at the beginning of the CURE), presumably in part due to encouragement by the instructors to seek out help when the students were feeling challenged (**Figure 5A**). Maladaptive coping strategies increased towards the later modules of the course, likely due to the students feeling pressure to finish final projects in all of the courses taken that semester as well as personal challenges that arose towards the end. The most frequently reported coping strategies over the entire course included reviewing the learning materials, demonstrating that the students found the learning materials helpful in completing the required analysis (**Figure 5B**).

**Figure 5.**
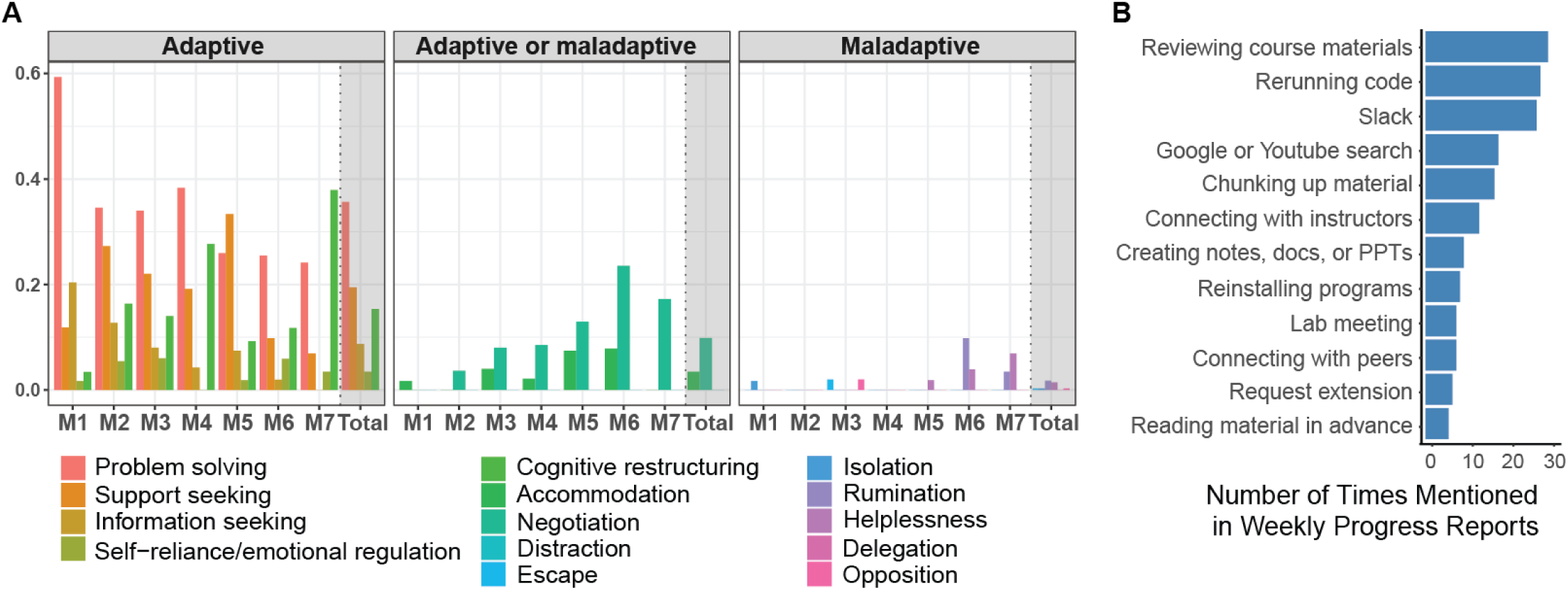
Coping themes categorized by coping type and frequency. (A) Coping strategies to overcome asynchronous course challenges self-reported in open-response progress reports each module, categorized as adaptive, maladaptive, or both depending on context. Adaptive themes include problem solving (red), support seeking (orange), information seeking (gold), self-reliance/emotional regulation (olive), and cognitive restructuring (green). Maladaptive themes include escape (light blue), isolation (blue), rumination (purple), helplessness (lilac), delegation (fuschia), and opposition (pink). (B) Most frequently reported student coping strategies from progress reports submitted throughout the course.

Identifying student-reported barriers helps make informed modifications while the CURE is being implemented. Instructors would meet independently each week to discuss the reported challenges and how to address them, with most of the solutions becoming the topic of the following week’s synchronous lab meeting discussion. These dynamic lab meetings created a virtual flipped classroom that derived lesson plans from student feedback and collaborated to find effective solutions specific content or code of concern. Lessons from analyzing coping strategies from progress reports can be used to provide guidance for students in future CUREs and may be helpful for instructors to figure out ways to encourage adaptive coping strategies in future CUREs.

### Effective student engagement in communication platforms

Engagement in communication platforms was quantified by exporting individual analytics from the Slack channel. At the time of this study, there were no grading options between Canvas and Slack available. There were no requirements for Slack engagement, but students were strongly encouraged to, at minimum, read the general class channel for updates and troubleshooting guidance. The ‘days active’ metric was used to evaluate how many days students were loading the channel page out of the total 51 days of the course duration whether or not the students offered any posts or replies. The ‘messages posted’ metric was used to quantify the number of messages a student posted in any of the course-specific channels, as an initial post or reply. To sort students by engagement, or frequency of message posting, average engagement was calculated by dividing the total conversation (set as 100%) by the number of students (n = 13). A posting threshold of average or higher was termed high engagement, from half of the average to the average was termed moderate engagement, and no posts to half of the average was termed low engagement. Interestingly, Slack activity showed a low positive correlation to normalized learning gains (R = 0.35, p-value = 0.2), but the percentage of Slack messages posted showed a high statistically significant correlation with normalized learning gains (R = 0.81, p-value = 0.0009) (**Figure 6**). This indicates that the students who were actively engaged in the course communications were able to build on their initial knowledge during the CURE, while those that watched the communications passively without participating were less likely to gain the knowledge being taught.

**Figure 6.**
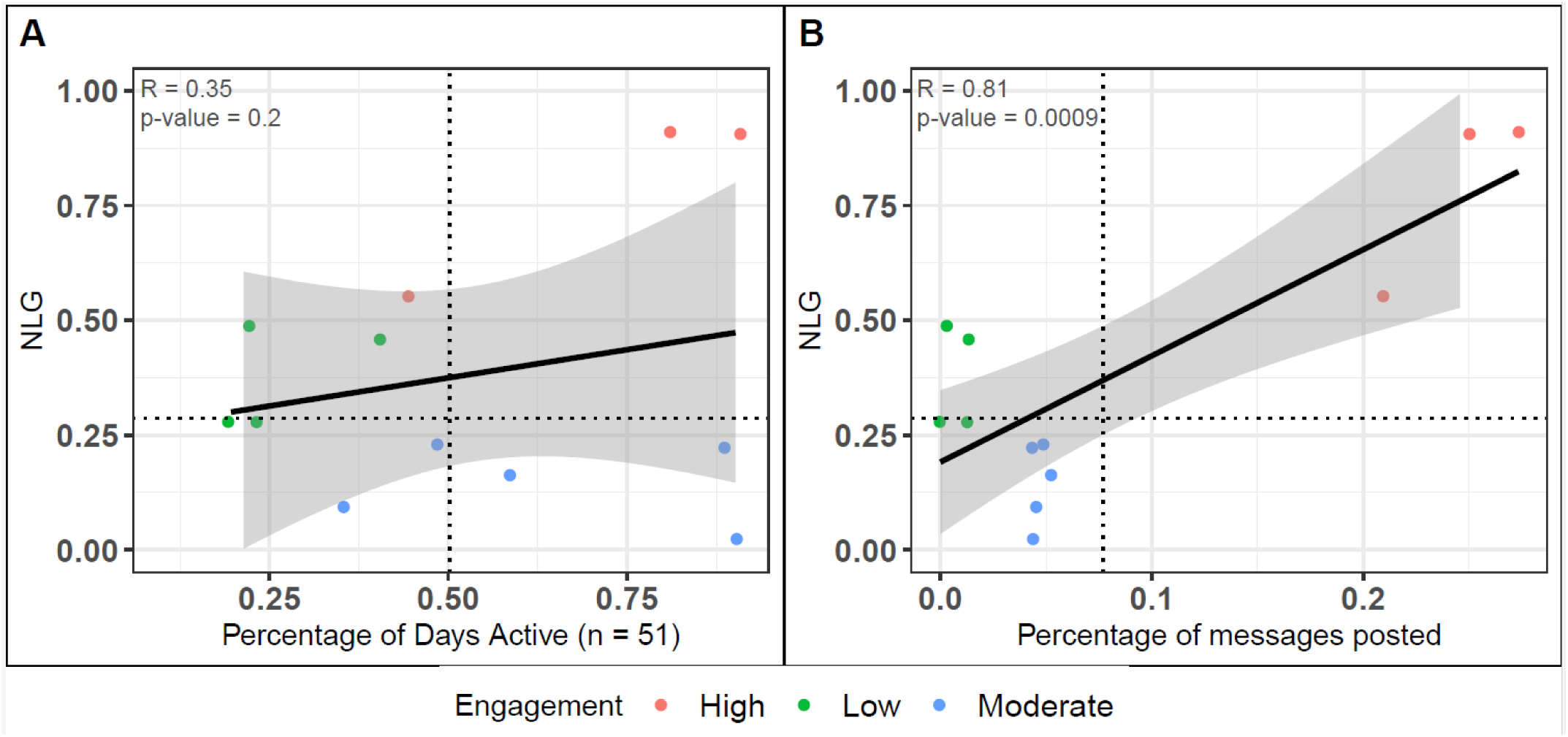
Normalized learning gains (NLG) by (A) Slack activity and (B) Slack messages posted. Student Slack engagement data was determined by pulling Slack analytics from course-specific channels from the day courses were made available until the end of the quarter. Engagement status was determined by dividing 100% of the conversation by the total number of students (n = 13 students) to determine if the theoretical average (7.69%) categorized as high (red, ≥ 7.69%), medium (blue, 3.81-7.68%), or low (green, ≤ 3.80). The horizontal, dotted line represents the mean class NLG (+28.63%). (A) Percentage of days active, or days of reading at least one channel or direct message, were determined by taking the individual’s number of days active and dividing it by the total number of days in the course (n = 51 days). The resulting percentages were plotted against student NLGs to determine correlation. Vertical dotted lines represent the mean number of days active from Slack analytics (25.6 days or 50.23%) (B) Percentage of messages posted conveys the number of messages students sent in the course-specific channel (excluding direct messages) divided by total messages received across the channel (n = 662 total posts). Vertical dotted lines indicate the theoretical average for engagement status, 7.69%.

## Conclusions and Future Directions

Upon completion of the genomics CURE, students were invited to continue to work on the research project to completion. The majority of the students (9 of 13) requested to continue on with the project by doing follow-up analysis or preparing the genomics results for publication. Follow-up studies focused on assessing the generalizability of the results from the CURE by testing the established protocols with other popular trimming tools and expanding the trimming parameter test space. The entire experiment will be repeated on a second set of placenta samples processed with RNA sequencing to test the reproducibility of the results. Once the results are at a level where they would be suitable for publication, students that remained active in the research after the CURE will be eligible for authorship and other students will be eligible for acknowledgement.

Furthermore, students sought out research and professional development opportunities outside of the CURE research project. The instructors are aware of 4 students who applied for Ph.D. programs shortly after completing the CURE, at least one of whom was not considering pursuing a Ph.D. until taking this course. Two students presented the research they did during the CURE at two undergraduate research symposiums the following semester. One was an online conference that used the Gathertown platform to present posters virtually. The other was a hybrid conference where a printed poster was hung next to a laptop computer from which the students presented through teleconferencing software. These two students placed in the top 3 bioinformatics presentations at the hybrid conference based on their mastery of the subject matter and exemplary communication skills. Both students expressed gratitude that they had an opportunity to present to a diverse audience and that they learned valuable lessons that they will use in graduate school.

The success of the students in the asynchronous genomics CURE has led to the development of future CURE research projects. As new research projects arise organically from discussions in the instruction team’s laboratory, suggestions are made for how those projects can be distributed to students and conducted asynchronously. Course development has involved tailoring learning objectives to suit the specific nature of the new CURE research project and custom learning materials required for the new project include a combination of newly and previously developed materials. This study demonstrates that students can be successful in online research experiences that incorporate channels for communication, bespoke and accessible learning materials, and open-response reports to monitor challenges and coping strategies. Additional scalable techniques for student assessment and communication will help to extend online research opportunities to many more students to come. Online asynchronous CUREs are a way to bring valuable research opportunities to many students who will go on to contribute to a more diverse and capable workforce to tackle an ever expanding set of genomics challenges.

## Acknowledgements

● London Skiles and Arizona State University EdPlus Instructional Design team for help getting learning materials on Canvas learning platform
● ASU Research Computing for help using the high performance computing cluster
● Funding sources: NSF-IUSE, OURS, NIH MIRA, HHMI Inclusive Excellence grant (S.E.B. is Co-PI)

## Author Contributions

● Melissa Wilson: Lead instructor and lead investigator
● Seema Plaisier: Course instructor, research project coordinator
● Danielle Alarid: Course instructor, analysis of student learning
● Katelyn M. Cooper and Sara E. Brownell: CURE experts and assessment experts
● Ken Buetow: Genomics expert

